# The cost of dual-task walking: Cognitive demands restrict visual search and gait planning

**DOI:** 10.1101/2025.11.16.688732

**Authors:** Y Russo, J Ye, SE Lamb, A Gear, W Lyon, J Qiu, M Banks, E Martin, Z Wang, M Alghamdi, P Leveridge, WR Young

## Abstract

Adaptive walking relies on proactive visual search to plan foot placement and maintain stability. This study examined how cognitive load and task complexity affect gaze behaviour and gait biomechanics during a precision target-stepping task in healthy young adults. We also quantified the frequency of cross-stepping during the experimental task.

Twenty-three participants (18–23 years) walked along an L-shaped pathway containing raised stepping targets under single-task (ST) and dual-task (DT) conditions. Targets had four different layouts to create high and low difficulty conditions. Eye movements were recorded using mobile eye-tracking, and gait kinematics were captured via motion capture.

Compared with ST, DT walking produced slower walking speeds, longer stance times, and reduced velocity between stepping targets, indicating a more inefficient gait strategy. Eye-tracking analyses revealed fewer and shorter fixations on task-relevant targets and a greater number of fixations directed toward task-irrelevant areas under DT conditions. In DT, saccadic amplitudes were reduced despite increased outside fixations, suggesting a breakdown in visual exploration between proximal and distal regions of the walkway. Cross-stepping was more frequent in ST conditions than in DT.

These findings indicate that increased cognitive load compromises proactive visual search, likely through working-memory and attentional limitations that disrupt feedforward gait planning. Contrary to expectation, cross-stepping occurred more often during ST than DT walking, suggesting that in this population cross-stepping may not be a maladaptive strategy. Overall, these results highlight the cognitive demands of adaptive walking even in young, low-risk individuals and underscore the importance of preserving visual–motor coordination under cognitive stress.

**Highlights:**

- Cognitive load during walking restricts proactive visual search and alters gait dynamics in healthy young adults.
- Dual-tasking led to slower walking speeds, longer stance times, and reduced step-to-step velocity.
- Eye-tracking showed fewer and shorter fixations on stepping targets, and more gaze directed toward task-irrelevant areas under dual tasking.
- Reduced proactive gaze under dual-task conditions likely reflects limitations of available cognitive resources rather than task-specific prioritisation.
- Findings highlight how cognitive demands compromise visual–motor coordination, supporting interventions that promote gait automaticity and reduce cognitive load.
- Cross-stepping was more frequent in ST than in DT conditions, suggesting that cross-stepping may not be maladaptive in healthy young people.

## Introduction

Navigating complex environments safely and efficiently requires us to continuously gather and integrate sensory information. Visual search is essential to adaptive gait as it provides the necessary information to precisely place the feet on areas deemed safe, avoid obstacles, and make ongoing adjustments ^1^. Many studies have reported characteristics of visual fixations and eye movements during adaptive gait as these provide an indication of planning efficiency^2,3^.

When completing complex precision-stepping tasks, young healthy adults typically demonstrate ‘proactive’ search (i.e., visual search of stepping constraints located beyond the nearest stepping constraint) ^4,5^. This proactive gaze strategy allows for effective sampling of the intended walking route so that walkers can make effective motor plans and transition efficiently between stepping constraints. In contrast, older adults, particularly those at higher risk of falling, often show reduced proactive visual search. It is thought that this behaviour might compromise walking safety, not only because of the reduced visual information acquired regarding subsequent constraints, but also because this restriction in visual search is associated with premature re-direction of visual attention away from an intended stepping location prior to the completion of the step. In other words, older ‘fallers’ look away from a stepping constraint before their step is completed in order to preview areas of the walkway where previous visual exploration of insufficient ^6^ Changes in visual search have been interpreted as maladaptive due to the association with increased stepping errors ^4,7^).

Walking is not purely automatic, it draws on attentional and executive functions ^8^, and studies incorporating dual-task (DT) paradigms have shown that when cognitive load increases, gait can be disrupted both in young and older adults, although it is most pronounced in the latter population ^9,10^. DT interference is particularly evident in high-risk populations and has been linked to greater fall risk ^11–14^. Beyond age-related decline, it is hypothesised that following negative adverse events (e.g. accidents, falls, injuries etc) individuals may start to consciously monitor and control their movements, especially under increased stress or task complexity, redirecting attention internally which generally results in reducing their ability to process external cues ^13–15^. Reinvestment is associated not only with compromised performance but also with impaired recollection of visuospatial features of the environment that participants have just traversed, limiting their ability to build or retain an accurate “spatial map” of the walkway. Conscious motor processing (CMP), as a manifestation of reinvestment, is cognitively demanding in its own right and has also been shown to restrict proactive visual search in both young and older adults ^5,16^. Together, these findings suggest that CMP may reduce the availability of attentional resources for previewing future stepping locations, thereby constraining efficient feedforward gait planning.

Previous research has evaluated the relationship between cognitive demands and visual search during adaptive gait during a relatively simple walking task ^17^. This study showed that DT in young adults did not reduce proactive visual search but instead induced visual disengagement from the intended walking path. Here, participants showed increased fixations on task-irrelevant areas outside the walkway; a phenomenon the authors referred to as “gazing into thin air”, alongside slower walking speeds and a higher frequency of stepping errors. Similarly, Feld & Plummer reported that participants performing a letter fluency DT directed more gaze toward task-irrelevant regions of a walking path compared to when walking without a cognitive load ^18^. Together, these findings challenge the assumption that cognitive load simply reduces proactive search of task-relevant areas of interest. Instead, they suggest a form of ‘structural interference’, where visual demands for gait planning compete with central executive resources required for the secondary task, resulting in active attempts to visually disengage from the intended walking path to prioritise the processing of a given concurrent cognitive task. While the studies described above have identified the potentially informative phenomenon of task-irrelevant ‘outside’ fixations in the context of gait, their paradigms were not sufficiently complex to evaluate the distribution of visual attention between proximal vs distal stepping constraints. Feld & Plummer did not integrate any precision stepping targets and Ellmers and colleagues created stepping targets using tape fixed to the ground with no requirement for specific foot orientation or significant consequences for stepping errors ^17,18^.

To our knowledge, there are no studies in the current literature describing the potential influence of increased cognitive demands on proactive visual search of multiple intended stepping locations.

In the current study we aimed to address these limitations by examining how healthy young adults adapt their gaze and gait in response to increased cognitive load and a manipulation of task complexity. Using a dual-task paradigm and a target-stepping layout with raised circular targets (i.e., to introducing features that would impact people balance in case of stepping error), we investigated how visual search strategies and gait biomechanics change under single-task and DT conditions. A key aim of this work was to examine how proactive gaze behaviour varies with environmental challenge and to explore whether these changes reflect cognitive resource constraints. We anticipate that DT conditions will provoke a greater number and frequency of outside fixations, even in more complex stepping conditions that increase demand for proactive visual search of target locations/orientations. We predict that failure to adequately fixate stepping targets in a proactive manner (i.e., fixating the target more than 2 steps prior to contact) will be associated with increased stepping errors ^5,6^. We also assessed the frequency of cross-stepping - the medial movement of one foot beyond the position of the other foot relative to the plane of the pelvis – which has been reported as a common cause of falls in older adults and is interpreted here as a risky stepping strategy, even in young healthy adults ^19^. We anticipate that failure to proactively visually search target locations (i.e., in DT conditions when outside fixations are potentially more prevalent) will be associated with increased incidence of either cross-steps or significant gait inefficiencies (i.e., prolonging stance time, stopping or making additional short steps to avoid cross-stepping) ^20^.

## Methods

### Study Design

This study incorporated a 2x2 within-subjects design (ST vs DT) x (Low vs High difficulty).

#### Participants

Twenty-three healthy young adults (age range: 18-23 years old) were recruited from the undergraduate student population at the University of Exeter. Participants were excluded if they had any neurological disorders, visual impairments, or musculoskeletal injuries that could affect gait or balance. Only individuals with corrected-to-normal vision using contact lenses were included; glasses were not permitted. All participants provided written informed consent prior to participation. The study was approved by the internal review board of the Department of Public Health and Sport Sciences, University of Exeter (528055).

#### Procedure

Participants completed a walking task along a predefined path consisting of two straight walkways connected at a 90-degree angle, forming an L-shaped path. Each straight walk measured ∼2.5 m (see Figure 1). The start and end points of the task were marked with tape placed approximately at hip width to standardise the initial stance. Participants were instructed to stand on the starting mark facing forward, preventing them from visually pre-scanning the walkway. As they walked, participants were required to step into circular gait orientation targets (raised ∼5 cm above ground level) positioned in predefined locations on the walkway. A blue target was located at the turning point between the two straight walkways, and two red targets were placed sequentially on the second straight walkway. Participants completed the walking task barefoot to minimize variability due to footwear and were instructed to walk at a comfortable pace, stepping into each target accurately with whichever foot felt natural.

**Figure 1.**
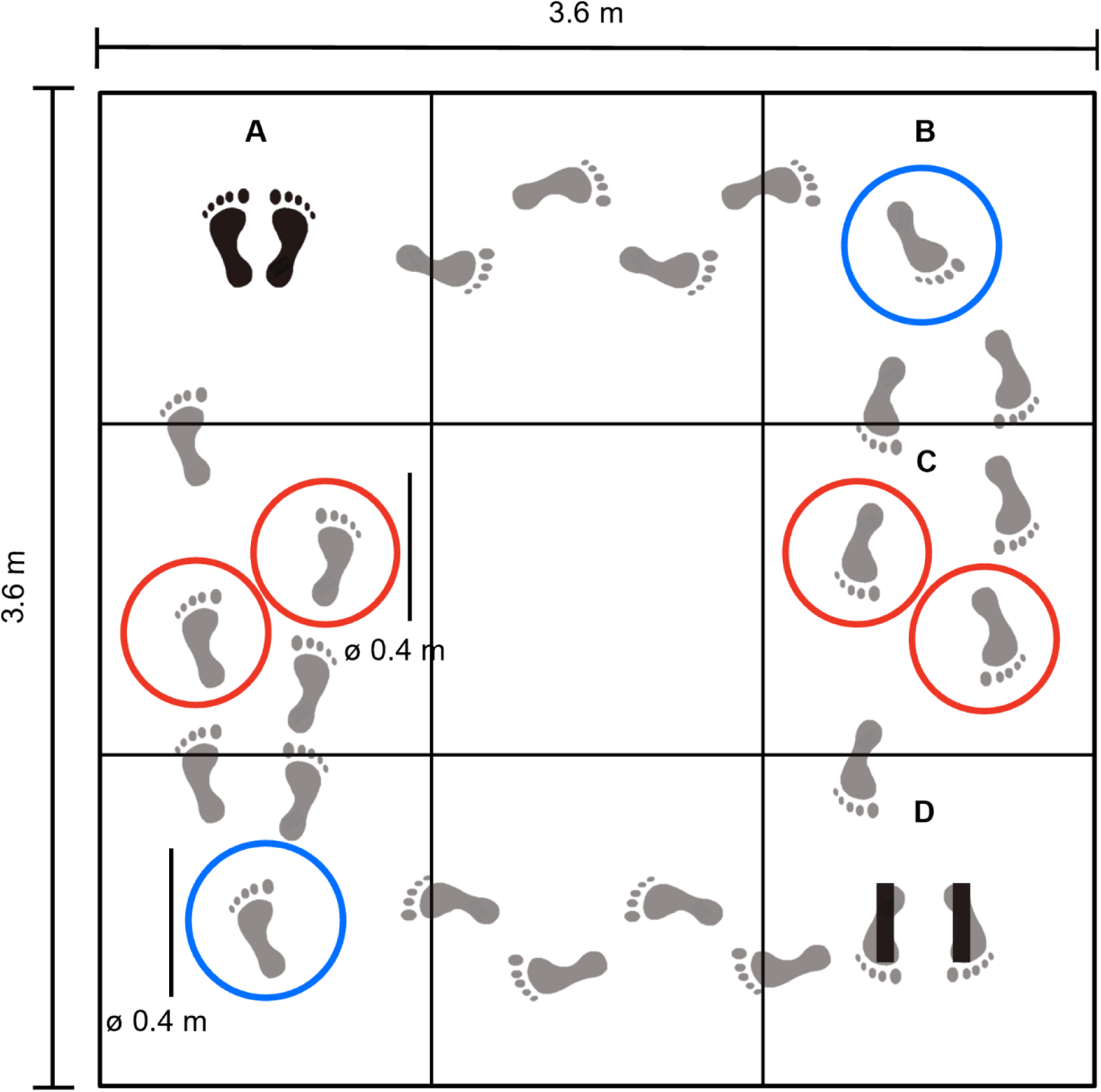
Layout of the experimental protocol. **A** marks the starting point of the walk. Participants began by facing away from the targets B and C to prevent them previewing of the walking path before the start. They then turned and walked straight from point A to point B. At **B**, the blue circle indicates where participants were required to step before turning towards the second straight walk. During this second walk, participants were instructed to step into the red circles (**C**) before stopping with their feet on the marks at **D**, with toes pointing towards the bottom edge of the platform. The exact arrangement of the circles at C varied from trial to trial (see Figure 2).

Four different target configurations were used, each representing a different difficulty level (Figure 2). Configurations A and C were considered as relatively simple, characterised by an everted initial stepping target (oriented outward), while configurations B and D were considered difficult, due to an inverted initial stepping target (oriented inward). The placement of the targets for the upcoming trial was adjusted at the very beginning of each walk, while participants were standing on the starting mark, looking forward.

**Figure 2.**
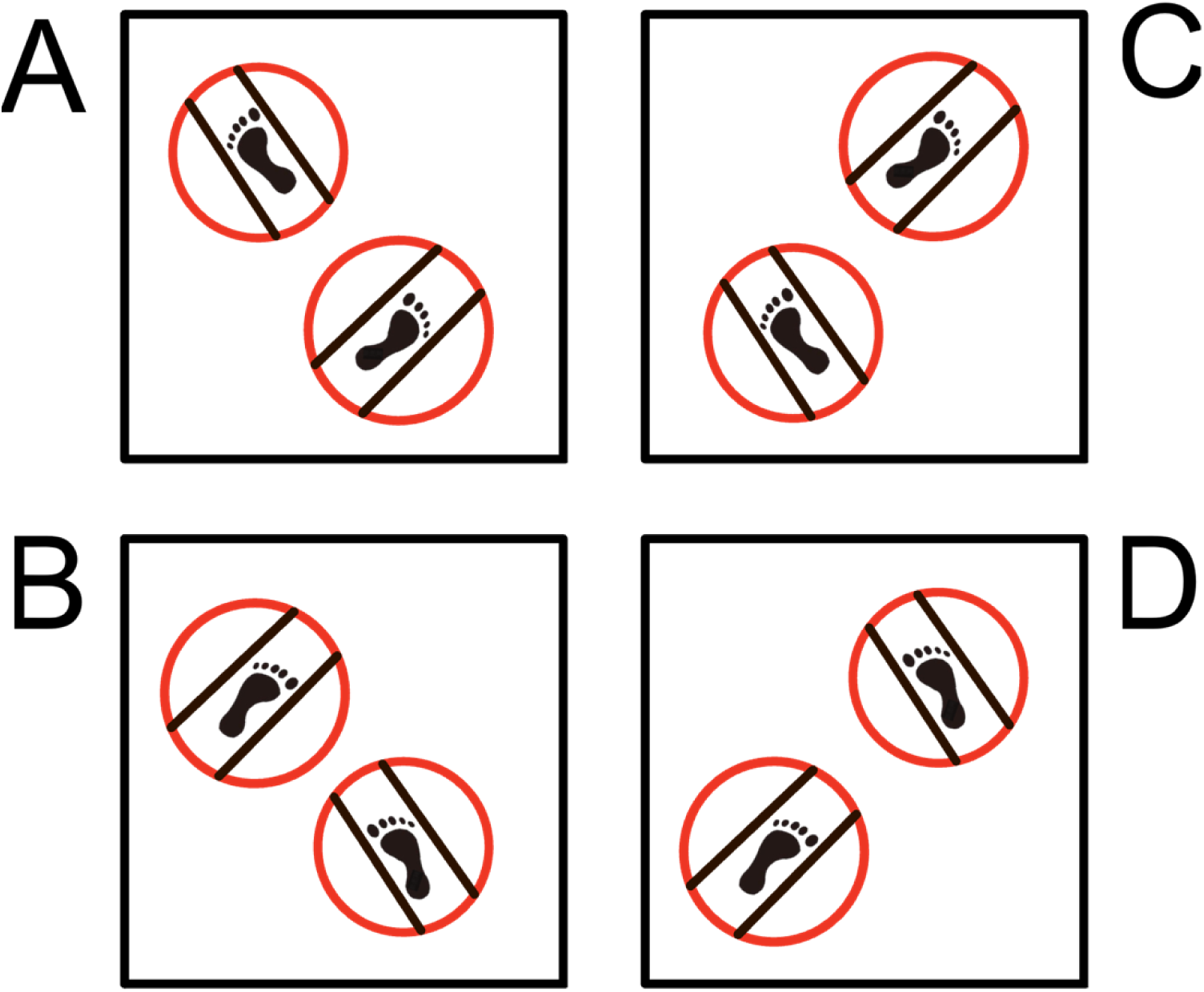
The four possible layouts of the red targets. Layouts **A** and **C** were considered simple because of their outward-oriented configurations. Layouts **B** and **D** were considered difficult due to the inward orientation and positioning of the feet within the targets.

By varying the position of the initial target, we were able to manipulate whether the natural footfall or walkers be conducive to stepping in the initial target on the ipsilateral side (i.e., right foot in right-sided target) or conducive for cross-stepping (i.e., the natural step length would lead the right foot to the left-side target, or vice-versa). To avoid either cross-stepping or gross stepping inefficiencies (i.e., stopping, making additional steps or rapid adjustments to step length), walkers needed to integrate these prospective requirements into their steps earlier in the approach to the targets shown in Figure 2.

Participants completed the walking task under two conditions: i) single-task (ST), when they had to complete only the walking task; ii) dual-task (DT): when they had to complete the walking task while simultaneously performing a cognitive task involving serial subtraction. Specifically, in the DT condition, participants were asked to subtract 7 multiple times (e.g., 203, 196, 189 etc.) starting from a number given by the experimenter, randomly selected between 185 and 210. Participants performed the subtraction aloud while walking.

Each participant performed both task conditions in separate, counterbalanced blocks: half of the participants began with the ST condition, and the other half with the DT. Within each block, participants performed multiple repetitions of the walking trial using different target configurations (i.e. simple and difficult). To reduce order effects, the layout and orientation of the gait targets were randomised and changed after every two laps. Each participant completed at least 3 trials in each target layout (Simple vs Difficult) and each condition (ST vs DT).

Before starting the experimental trials, participants were allowed to familiarise themselves with the walking task and DT condition, to ensure they understood the procedure and could perform the task adequately.

#### Instruments

Kinematic data were collected using a 19-camera motion capture system (PrimeX 13, OptiTrack, USA) recording at 100 Hz. Each participant wore 12 retro-reflective markers placed on three anatomical landmarks of each foot (heel, lateral malleolus, and head of the first metatarsal), the lateral femoral condyle of each knee, and both the anterior and posterior superior iliac spines. Eye movements were recorded at 100 Hz using an eye-tracking system (Tobii Pro Glasses 2, Sweden), which also captured first-person video of the trials at 25 Hz (1920×1080 pixels). Additionally, a video camera (HDR-CX405, Sony, Japan) recorded the trials at 25 Hz to assist with the identification of cross-steps and other gait events.

#### Data Analysis and Outcome Variables

Video recordings were reviewed by experimenters (ZW and JQ), who annotated each trial to determine which foot landed in which target and whether a gross stepping error or cross-step occurred.

Kinematic data were labelled, visually inspected, and analysed semi-automatically using custom scripts developed in MATLAB (R2023a, MathWorks, Natick, MA, USA). All kinematic data were low-pass filtered with a fourth-order Butterworth filter using a 15 Hz cut-off frequency (Rum et al., 2023). Individual trials were identified based on spatial thresholds within the motion capture coordinate system. Specifically, a trial was defined as beginning with the participant’s first step motion and ending when the pelvis markers crossed a threshold placed approximately 1 metre in beyond the second red target (see Figure 1).

For each trial, steps were identified based on heel strikes and toe-offs. A heel strike was defined as the minimum vertical position of the heel marker following a local maximum, while toe-off was identified as the minimum vertical position of the toe marker preceding a local maximum.

As target locations remained consistent across trials, their positions in each configuration (A, B, C, D) were approximated graphically in a MATLAB GUI. An experimenter (YR) manually identified specific steps of interest in each trial: the step before the blue target, the step into the blue target, the step into the first red target, and the step into the second red target. These steps were used to compute the following biomechanical outcomes: i) *stance duration* (time from heel strike to toe-off) for the step before the blue target, the step into the blue target, and the step into the first red target; ii) *average velocity* between the blue target and the first red target, calculated as the time between heel strikes on the blue and first red targets; iii) *step interval* time between the first and second red targets, defined as the time between their respective heel strikes. The annotated video data were used to cross-check the accuracy of the detected steps. Additionally, *task duration* was computed as the time from trial start to end, whereas *average walking speed* was calculated by dividing the walking path length (5 m) by task duration.

Eye-tracking data were manually labelled frame-by-frame in Tobii Pro Lab (v1.217) by the experimenters (JY, AG, WL, JQ, MB, EM, MA). Five areas of interest (AOIs) were defined: area before the blue target, blue target, gap, first red target, and second red target. For the analysis, each trial was segmented into two phases: (i) before stepping into the blue target and (ii) after stepping into the blue target. This distinction allowed us to differentiate between instances in which the red targets and the gap were viewed as part of proactive visual search (i.e. planning future steps) versus proximal visual search (i.e. guiding immediate foot placement).

#### Statistical Analysis

Data normality was assessed using the Shapiro–Wilk test. Effect sizes were reported as partial eta squared (η²). A 2 × 2 repeated measures analysis of variance (RM-ANOVA) was conducted to compare biomechanical and eye-tracking variables across cognitive conditions and difficulty levels. All statistical analyses were performed using SPSS (version 28, IBM Corp., Armonk, NY, USA). The significance level was set at α = 0.05 for all tests.

## Results

### Participants

Of the 23 participants recruited, 6 were excluded from the final analysis. Of these, 2/6 participants did not complete the ST condition, and 4/6 had fewer than two valid trials in the DT condition with the difficult target layout. Additionally, 4 participants were excluded from the biomechanical analysis (but not the eye tracking one) due to issues with motion capture recordings. One further participant (PT22) was excluded from biomechanical analyses involving the second red target due to marker visibility issues in the DT condition.

As a result, the final sample included 17 participants for the eye-tracking analysis and 13 participants for the biomechanical analysis. On average, the 17 participants included in the eye-tracking analysis completed 12 ST walks (±5 trials) and 11 DT walks (±5 trials).

### Main Outcomes

#### Biomechanical Outcomes

Cognitive task had a significant effect on several biomechanical variables (see Table1). Trial duration was significantly longer during DT compared ST walking. Consistently, average walking speed was significantly reduced in DT trials, compared to ST.

Cognitive task also influenced gait parameters associated with specific targets (Table 1). Stance time on the initial blue target was significantly longer in DT than in ST. Similarly, gait velocity between the blue and the first red target was significantly higher in ST compared to DT. Stance time on the first red target was also significantly longer in DT than ST. Finally, the time taken to step between the first and second red targets was significantly greater in DT than ST (Table1).

**Table 1.**
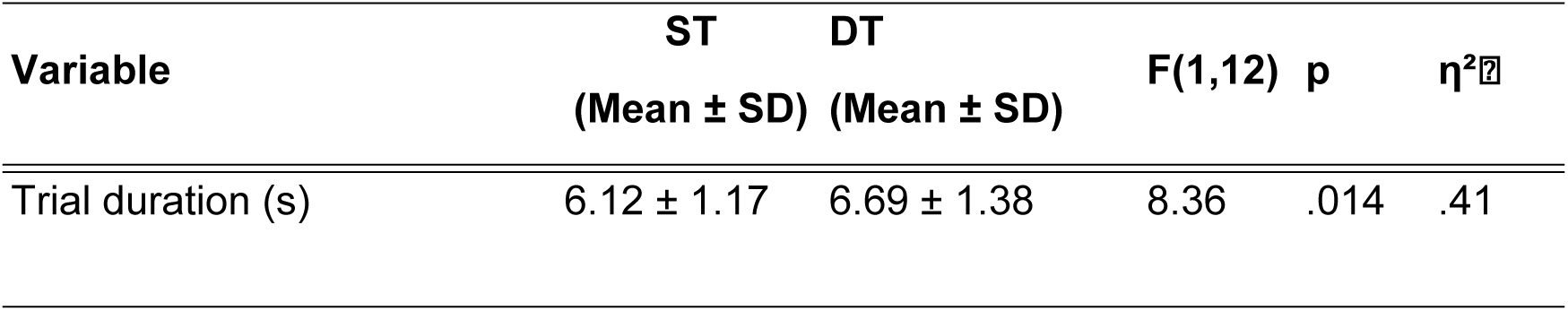
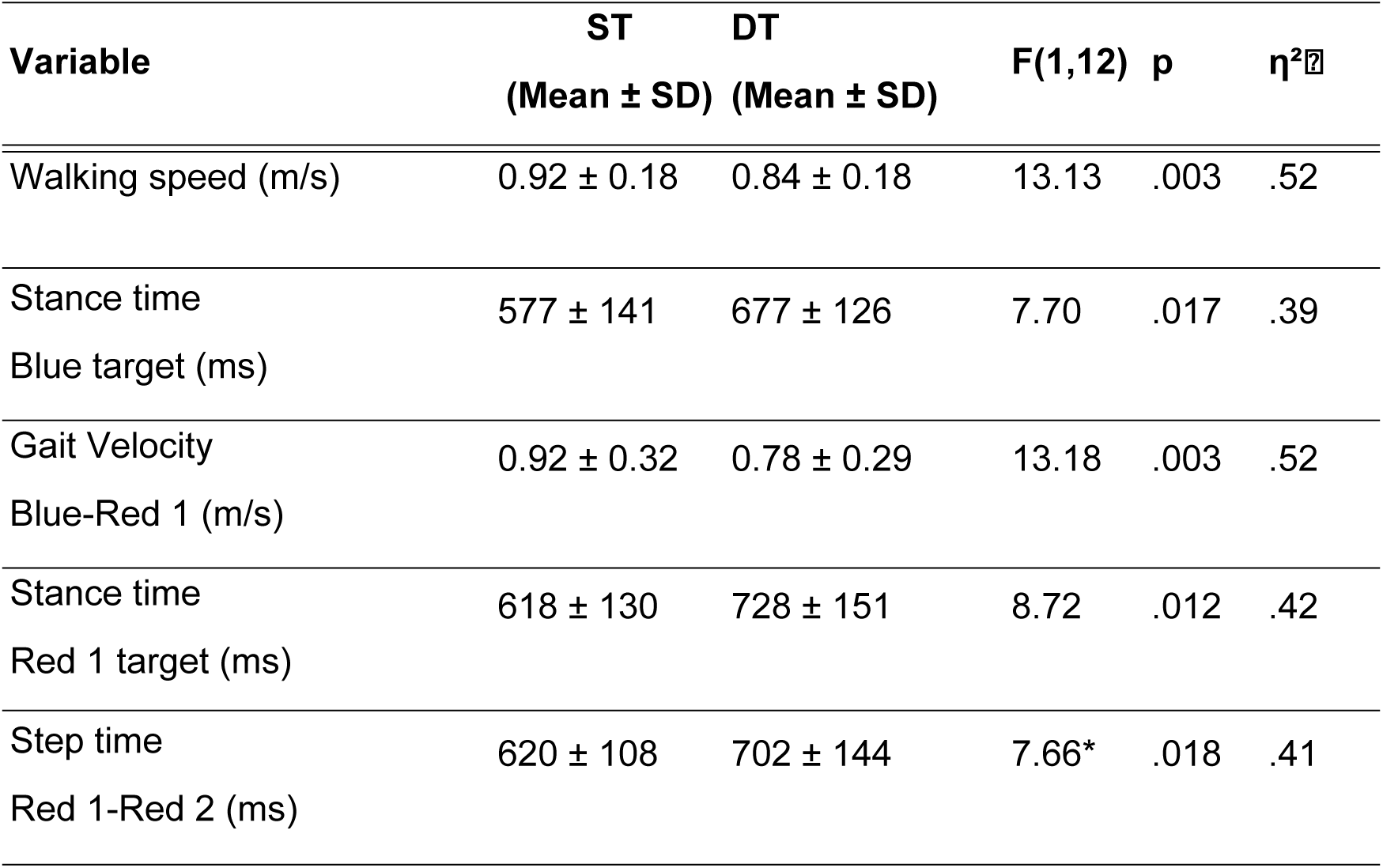
reports significant differences between conditions for biomechanical outcomes. Mean (±SD) gait parameters are shown for single-task (ST) and dual-task (DT) walking conditions. Effect sizes are reported as partial eta squared (η²). *Please note that step time degrees of freedom for step time Red1-Red2 are (1, 11).

| Variable | ST<br>(Mean ± SD) | DT<br>(Mean ± SD) | F(1,12) | p | $\eta^2$ |
| --- | --- | --- | --- | --- | --- |
| Trial duration (s) | 6.12 ± 1.17 | 6.69 ± 1.38 | 8.36 | .014 | .41 |

| Variable | ST<br>(Mean ± SD) | DT<br>(Mean ± SD) | F(1,12) | p | η<sup>2</sup> |
| --- | --- | --- | --- | --- | --- |
| Walking speed (m/s) | 0.92 ± 0.18 | 0.84 ± 0.18 | 13.13 | .003 | .52 |
| Stance time<br>Blue target (ms) | 577 ± 141 | 677 ± 126 | 7.70 | .017 | .39 |
| Gait Velocity<br>Blue-Red 1 (m/s) | 0.92 ± 0.32 | 0.78 ± 0.29 | 13.18 | .003 | .52 |
| Stance time<br>Red 1 target (ms) | 618 ± 130 | 728 ± 151 | 8.72 | .012 | .42 |
| Step time<br>Red 1-Red 2 (ms) | 620 ± 108 | 702 ± 144 | 7.66* | .018 | .41 |

No significant differences were found in biomechanical outcomes for task difficulty nor significant task condition x motor difficulty interactions.

#### Eye-Tracking Outcomes

When examining gaze behaviour on the first red target in in the period when approaching the initial blue target (i.e., the period when fixations on the red targets would be categorised as proactive), both the number and duration of fixations were significantly reduced in DT (Table 2 and Figure 3, top panels). Specifically, DT trials showed a lower number of fixations compared to ST trials, and shorter average fixation durations.

**Figure 3.**
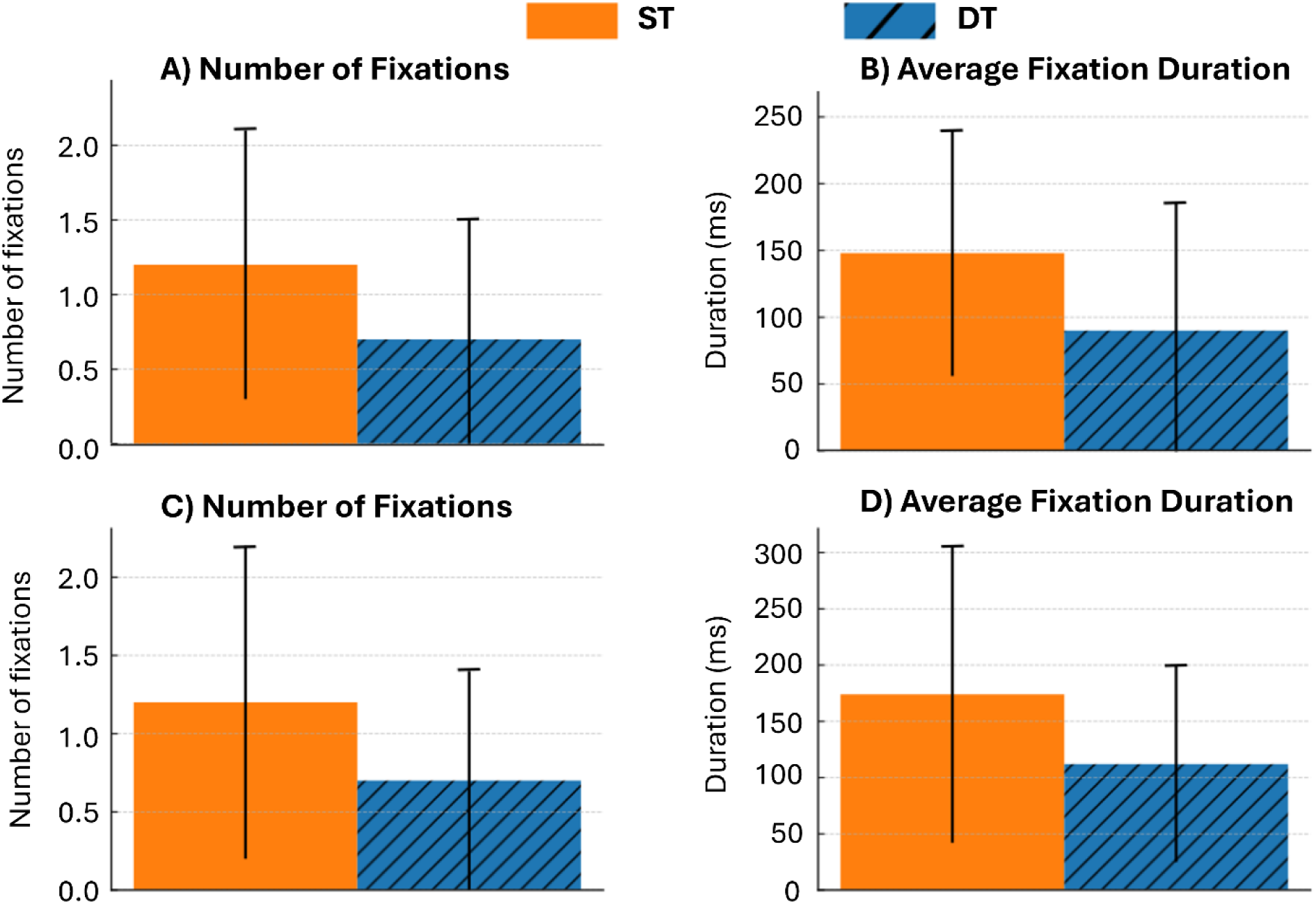
Summary of fixation behaviour (i.e., number of fixations and average fixation duration) on the first red target. Panels A and B show fixation characteristics when the first red target was viewed while approaching the blue target (i.e., proactive visual search). Panels C and D show fixation characteristics on the first red target after foot contact on the blue target, that is, when the first red target acted as a proximal obstacle along the pathway. ST = single task; DT = dual task; ms = milliseconds.

**Table 2.** reports significant differences between conditions for eye-tracking outcomes. Mean (±SD) gait parameters are shown for single-task (ST) and dual-task (DT) walking conditions. Effect sizes are reported as partial eta squared (η²).

| Variable | ST Mean ± SD | DT Mean ± SD | F(1,16) | p | $\eta^2$ |
| --- | --- | --- | --- | --- | --- |
| Fixations on red target<br>(approach phase, number) | 1.2 ± 0.9 | 0.7 ± 0.8 | 10.57 | .005 | .40 |
| Fixation duration on red<br>(approach phase, ms) | 148 ± 92 | 90 ± 96 | 5.88 | .028 | .28 |
| Fixations on red<br>(proximal target, number) | 1.2 ± 1.0 | 0.7 ± 0.7 | 13.61 | .002 | .46 |
| Fixation duration on red<br>(proximal target, ms) | 174 ± 132 | 112 ± 87 | 10.76 | .005 | .40 |
| Number of Fixations<br>on outside area (number) | 3.43 ± 2.1 | 6.25 ± 3.89 | 15.29 | .001 | .49 |
| Saccade amplitude ($^\circ$) | 10.14 ± 1.76 | 9.07 ± 2.02 | 8.14 | .012 | .34 |

A similar pattern was observed when the first red target was viewed in the period between foot contact in the first blue target and foot contact in the initial red target (Table 2 and Figure 3, bottom panels). The number of fixations was significantly smaller in DT than in ST. The average duration of fixation on this target was also shorter in DT than ST.

Gaze behaviour on the outside area also differed between cognitive tasks. Participants made significantly more fixations on outside areas during DT trials compared to ST (Table 2). However, there were no significant differences in average fixation duration on the outside area between motor difficulty levels.

Eye-tracking analyses revealed that cognitive tasks significantly affected the average amplitude of saccades, participants made larger saccades in ST than in DT trials.

Notably, a significant condition x motor difficulty interaction emerged for average fixation duration on the outside area, *F*(1,16) = 4.609, *p* = 0.047, η² = 0.224. In ST trials, fixation duration on the outside area increased from the easy (305 ms ±44) to the hard layout (404 ms ±74), while in DT trials, fixation durations remained relatively unchanged (easy: 324 ms ±42; hard: 303 ms ±45). No other significant effects of condition or difficulty were observed.

### Secondary Outcomes

#### Descriptive Analysis of Cross-Steps

A total of 57 cross-steps were observed across the 17 participants included in the analysis. Of these, 13 participants exhibited cross-stepping behaviour when stepping onto the red targets, with a range of 1 to 11 cross-steps per participant (median = 3; interquartile range = 8.5). Notably, cross-steps occurred more frequently during single-task trials (n = 43) than during dual-task trials (n = 14). Due to the heterogeneous distribution of cross-steps across participants and across conditions, inferential statistical comparisons between cross-steps and standard steps were not performed. Descriptive biomechanical characteristics of cross-steps and non-cross steps in ST and DT conditions are presented in Table 3.

**Table 3.** provides an overview of the biomechanical outcome measures, divided according to task type (single vs. dual task) and the presence of cross-stepping. ST = single task; DT = dual task.

|  | Trial<br>Duration<br>(s) | Average<br>Velocity<br>(m/s) | Stance<br>Before<br>Blue<br>(ms) | Stance<br>in Blue<br>(ms) | Velocity<br>between<br>blue and<br>red (m/s) | Stance<br>in 1<sup>st</sup><br>Red<br>(ms) | Stance<br>in 2<sup>nd</sup><br>Red<br>(ms) |
| --- | --- | --- | --- | --- | --- | --- | --- |
| ST<br>Cross-Step | 5.88±1.25 | .93±.20 | 639±127 | 563±154 | .92±.29 | 617±205 | 640±170 |
| ST Non-Cross-Step | 6.59±2.24 | .81±.17 | 664±133 | 643±145 | .92±.30 | 666±224 | 644±153 |
| DT<br>Cross-Step | 6.07±1.24 | .86±.17 | 699±174 | 694±238 | .73±.20 | 679±174 | 651±257 |
| DT Non-Cross-Step | 6.87±1.63 | .77±.16 | 690±145 | 693±163 | .79±.33 | 748±154 | 725±195 |

## Discussion

This study investigated how cognitive load and task difficulty influence visual search behaviour and gait biomechanics in healthy young adults during a target-stepping task. Our results show that dual-task significantly restricted gaze behaviour and influenced locomotor performance, providing new insights on the effects of cognitive demands of adaptive gait in young populations.

### Proactive Visual Search is Disrupted Under Dual-Task Conditions

Participants walked more slowly and exhibited longer stance times when engaged in a DT compared to a ST, consistently with the broader literature showing that cognitive load impairs motor performance ^8,10,17^. Reduced velocity between stepping targets and prolonged stance times suggest that participants adopted a more cautious gait strategy under cognitive load, possibly a compensation for disruptions in feedforward planning and therefore reduced visual-motor coordination. Eye-tracking data provided further support for this hypothesis. Under DT conditions, participants fixated less frequently and for shorter durations on task-relevant areas, especially the first red target, and directed their gaze more towards task-irrelevant outside areas. This shift in gaze behaviour reflects a breakdown in proactive visual search, which is critical for feedforward motor planning. These findings corroborate previous work by Ellmers, who observed that cognitive load induced “gazing into thin air”, a form of active visual disengagement from the walking task ^17^. In this work, the experimenters observed similar restrictions in visual search, interpreted by the authors as a by-product of the cognitive demands associated with these manipulations ^17^. However, no prior study had demonstrated a clear relationship between increased cognitive demands and restricted proactive visual search, until now. The current findings extend this literature by revealing how dual tasking disrupts proactive search. Specifically, we show that cognitive DT reduces saccadic amplitude even when walkers are making larger saccades to fixate outside area, suggesting a possible breakdown in visual exploration between proximal and distal areas of the walkway. Therefore, this behaviour could represent an adaptive attempt to stabilise gaze under higher cognitive load. However, we observed that DT only increased the number of outside fixations, and no concurrent differences were observed in fixation durations; changes that one would expect to observe if stabilisation of visual scene was being prioritised. Altogether, these results suggest that reduced cognitive capacity under DT conditions compromises visual search functions. This interpretation is reinforced by the reduction in fixations on the first red target following blue target contact in the DT condition. Whereas previous studies manipulating CMP found increased fixations on proximal constraints ^16^ - reflecting prioritisation of nearby task-relevant information - the present results show the opposite pattern. In this context, CMP might be considered as more task-specific and therefore responsible for driving proximal fixations relevant to control of ongoing/upcoming movements. In contrast, processes relating to cognitive DT are irrelevant to the motor task. Here, the absence of proximal fixations (associated in other studies with CMP) combined with a clear reduction in proactive search implicates search functions and processes relating to working memory as the likely cause of reduced proactive search. This is potentially in contrast with CMP-related changes as reduced search in those may have occurred due to the prioritisation of proximal fixations rather than reduced cognitive capacity. Our findings therefore support the view that anxiety - and CMP-related changes in visual behaviour may, at least in part, stem from the heightened cognitive demands associated with these manipulations. This interpretation highlights the importance of fall-prevention interventions that promote gait automaticity and reduce the cognitive load of walking, thereby preserving proactive visual search and supporting safer locomotion.

However, while increased cognitive demands appear to constrain proactive visual search, they do not seem to fully account for the more pronounced proximal fixations and reliance on online control. In this context, researchers and health professionals should consider the extent to which CMP may also be adaptive (i.e., beneficial in ensuring safety). Consequently, rather than aiming solely to restore behaviours typical of young, healthy adults, researchers and clinicians should consider the extent to which CMP may compensate for age-related decline. Future work should identify the point at which CMP transitions from adaptive to maladaptive, thereby informing rehabilitation strategies that preserve effective compensatory mechanisms without compromising safety. We argue that this is necessary in order to establish clear targets for rehabilitation that do not compromise effective compensatory strategies.

Previous work has also shown that restrictions in visual search are associated not only with increased stepping errors but also with longer stance phase durations around precision steps^5^. The present findings further support this suggested link. A failure to proactively acquire visuospatial information about upcoming target locations and orientations is likely to compromise feedforward movement planning during the approach to the targets. As a result, walkers would slow down, particularly during the critical phases of gait where planning of future steps and propulsion of the centre of mass are carried out. However, our findings - and those of previous studies ^17^ - suggest that participants may still attempt to maintain overall walking velocity despite these visuomotor constraints. In such cases, walkers appear to compensate by prolonging critical stance phases (e.g. stance in the blue target) rather than reducing speed substantially, a strategy that may preserve forward momentum but at the cost of reduced stepping accuracy (see also studies on anxiety and cognitive-motor interference ^6,22^).

### Outside Fixations May be Indicative of Visuomotor ‘Spare Capacity’

Participants produced more outside fixations during the DT condition. In accordance with previous reports of this behaviour ^17^, this finding appears to represent a structural interference between the motor task of walking and concurrent cognitive task. However, the current study helps to progress our understanding of this phenomenon due to the additional manipulation of motor task complexity. One might expect that increasing demands of the motor task – thereby increasing the importance of proactively visually searching the red targets - would restrict visual disengagement from the walkway. Our results showed no significant effect of motor task difficulty on visual search behaviour directed toward the red targets. However, the duration of outside fixations increased in the more complex motor task, but only during ST conditions. This finding directly contradicts our initial predictions and highlights the need to consider the origins and functions of outside fixations. Our results support the notion that high cognitive demands can be sufficient to induce outside fixations by virtue of an active disengagement that is presumably intended to avoid visual distractions associated with movement planning while prioritising the cognitive task. However, the same rationale cannot be applied to the ST condition. Here, the longer outside fixation durations are unlikely to be driven by any increased cognitive demands of processing information relating to the more complex target locations. We speculate that, upon recognising a more complex arrangement of targets, walkers developed an implicit understanding that there will be greater demands for motor planning in the step preceding the red targets. Once the constraints of the upcoming task are understood, the walker is at liberty to visually search the path or temporarily disengage without cost to the motor task. In this sense, we recommend that outside fixations should be considered as an expression of ‘spare capacity’ in the visuomotor system.

It is important to note that the current results suggest that in more difficult gait tasks, cognitive DT is not sufficient to drive outside fixations. In other words, due to the added requirements for movement planning, there is less capacity to disengage the gaze from the walkway. However, in the ST high difficulty tasks we observed more outside fixations. This contradicts the notion that increased requirements for movement planning are negatively associated with outside fixations and instead implies that those requirements might predispose people to visually disengage. Previous reports of outside fixations interpret it as a strategy to temporarily prioritise the cognitive task by avoiding task-relevant fixations ^17^. In the current task, the adaptive gait task was far more challenging compared to previous studies, especially around the red targets in the hard condition. This raises the possibility that, upon recognising the harder arrangement of red targets, walkers engaged in a strategy to retain this information, and this emerged as visual disengagement from task-relevant areas during periods of the task where precision stepping was not required. These mechanisms are likely to also have been present in DT conditions. However, we speculate the greater number of outside fixations observed in DT conditions is likely to have masked this effect. This is likely to have been particularly true in the easy target condition, as outside fixations is already shown to be increased by DT in easier gait tasks ^17,23^.

### Cross-Stepping as an Adaptive and Context-Dependent Strategy

Contrary to our initial hypothesis, we observed a higher frequency of cross-steps in the ST condition compared to the DT condition. Cross-stepping is often interpreted as a challenging compensatory strategy, typically associated with spatial or timing constraints and thought to reflect inefficiencies in gait planning. However, in this context, we propose an alternative explanation. in ST trials, participants were likely able to allocate more cognitive resources to the motor task, allowing them to explore a broader range of stepping strategies, including cross-steps, without compromising dynamic balance. Although the raised targets required increased precision for foot placement, they may not have represented a substantial threat to balance in this population. As such, cross-stepping may have been perceived as a safe and efficient solution under ST conditions, as suggested by the observed velocities reached during the different tasks (Table 3). In contrast, under DT conditions, the added cognitive load may have reduced participants’ confidence in adopting a biomechanically more challenging strategy. This could have led to a more conservative approach, favouring the adoption of normal stepping strategies that required less attentional regulation and minimised perceived risk. These findings partly challenge the notion that cross-stepping is inherently maladaptive and suggest that the selection of stepping strategy is context and population dependent and influenced by perceived motor capacity. Future studies should investigate the functional role and perceived risk of cross-stepping across different populations and environmental challenges, particularly in older adults or people with impaired balance, to determine whether similar patterns emerge under varying cognitive and physical constraints.

## Limitations

Several limitations should be acknowledged. First, the sample was limited to healthy young adults, and findings may not be generalised to older or clinical populations who exhibit different gait and cognitive abilities. Second, although the study used a well-controlled target-stepping paradigm, real-world walking environments are more variable and unpredictable. Finally, due to the uneven and limited distribution of cross-step events, we were unable to perform inferential comparisons between cross- and standard stepping behaviours.

## Conclusions

This study demonstrates that dual-tasking compromises both visual search and gait performance in healthy young adults, particularly by reducing proactive gaze behaviour and slowing stepping dynamics. These results highlight the cognitive demands of adaptive walking, even in low-risk populations, and underscore the importance of visual attention in locomotor planning. By identifying how cognitive load disrupts the spatial and temporal allocation of gaze during walking, this work contributes to a deeper understanding of attentional control during movement and informs the development of interventions aimed at enhancing safety in more complex and cognitively challenging environments.

## Author Contribution

Conception and design – YR, JY & WY. Acquisition of data - YR, JY, GA, LW, PL, ZW and WY. Data curation – YR, JY, AG, LW, JQ, MB, EM, ZW, MA. Formal analysis - YR, JY. Writing original draft - YR & WY. Writing review & editing - All the authors. Final approval of the completed article - All the authors.

## Acknowledgments

We would like to thank the participants who took part in the study.

## References

1. Land M, Tatler B. Looking and Acting: Vision and Eye Movements in Natural Behaviour. First. Oxford: OUP Oxford, 2009.

2. Hardiess G, Hansmann-Roth S, Mallot HA. Gaze movements and spatial working memory in collision avoidance: A traffic intersection task. Front Behav Neurosci. Epub ahead of print 6 June 2013. DOI: 10.3389/fnbeh.2013.00062.

3. Foulsham T. Eye movements and their functions in everyday tasks. In: Eye (Basingstoke). Nature Publishing Group, 2015, pp. 196–199.

4. Young WR, Wing AM, Hollands MA. Influences of state anxiety on gaze behavior and stepping accuracy in older adults during adaptive locomotion. Journals of Gerontology - Series B Psychological Sciences and Social Sciences 2012; 67 B: 43–51.

5. Ellmers TJ, Young WR. The influence of anxiety and attentional focus on visual search during adaptive gait. J Exp Psychol Hum Percept Perform 2019; 45: 697–714.

6. Young WR, Williams MA. How fear of falling can increase fall-risk in older adults: Applying psychological theory to practical observations. Gait and Posture 2015; 41: 7–12.

7. Young WR, Hollands MA. Can telling older adults where to look reduce falls? Evidence for a causal link between inappropriate visual sampling and suboptimal stepping performance. Exp Brain Res 2010; 204: 103–113.

8. Yogev-Seligmann G, Hausdorff JM, Giladi N. The role of executive function and attention in gait. Movement Disorders 2008; 23: 329–342.

9. Stavrinos D, Byington KW, Schwebel DC. Distracted walking: Cell phones increase injury risk for college pedestrians. J Safety Res 2011; 42: 101–107.

10. Agmon M, Belza B, Nguyen HQ, et al. A systematic review of interventions conducted in clinical or community settings to improve dual-task postural control in older adults. Clinical Interventions in Aging 2014; 9: 477–492.

11. Beauchet O, Annweiler C, Dubost V, et al. Stops walking when talking: A predictor of falls in older adults? European Journal of Neurology 2009; 16: 786–795.

12. Nagamatsu LS, Voss M, Neider MB, et al. Increased Cognitive Load Leads to Impaired Mobility Decisions in Seniors at Risk for Falls. Psychol Aging 2011; 26: 253–259.

13. Jackson RC, Ashford KJ, Norsworthy G. Attentional Focus, Dispositional Reinvestment, and Skilled Motor Performance Under Pressure. 2006.

14. Masters R, Maxwell J. The theory of reinvestment. Int Rev Sport Exerc Psychol 2008; 1: 160–183.

15. Young WR, Olonilua M, Masters RSW, et al. Examining links between anxiety, reinvestment and walking when talking by older adults during adaptive gait. Exp Brain Res 2016; 234: 161–172.

16. Ellmers TJ, Young WR. Conscious motor control impairs attentional processing efficiency during precision stepping. Gait Posture 2018; 63: 58–62.

17. Ellmers TJ, Cocks AJ, Doumas M, et al. Gazing into thin air: The dual-task costs of movement planning and execution during adaptive gait. PLoS One; 11. Epub ahead of print 1 November 2016. DOI: 10.1371/journal.pone.0166063.

18. Feld JA, Plummer P. Visual scanning behavior during distracted walking in healthy young adults. Gait Posture 2019; 67: 219–223.

19. Robinovitch SN, Feldman F, Yang Y, et al. Video capture of the circumstances of falls in elderly people residing in long-term care: An observational study. The Lancet 2013; 381: 47–54.

20. Lyon IN, Day BL. Predictive control of body mass trajectory in a two-step sequence. Exp Brain Res 2005; 161: 193–200.

21. Rum L, Russo Y, Vannozzi G, et al. “Posture first”: Interaction between posture and locomotion in people with low back pain during unexpectedly cued modification of gait initiation motor command. Hum Mov Sci; 89. Epub ahead of print 1 June 2023. DOI: 10.1016/j.humov.2023.103094.

22. Ellmers TJ, Wilson MR, Kal EC, et al. The perceived control model of falling: developing a unified framework to understand and assess maladaptive fear of falling. Age and Ageing; 52. Epub ahead of print 1 July 2023. DOI: 10.1093/ageing/afad093.

23. Cocks AJ, Young WR, Ellmers TJ. Reduced cognitive resources induces risky stepping strategies in older adults. Gait Posture. Epub ahead of print 2025. DOI: 10.1016/j.gaitpost.2025.109989.

